# Transcriptomic analysis of DNA damage response in zebrafish embryos under simulated microgravity

**DOI:** 10.1101/2021.09.30.462627

**Authors:** Subhrajit Barua, Aleksey Komissarov, Harpreet Kaur, Elia Brodsky

**Affiliations:** SCAMT, ITMO University, St. Petersburg, Russia; Pine Biotech, New Orleans, Louisiana, USA

**Keywords:** astrobiology, spaceflight, simulated microgravity, radiation, bioinformatics, space omics, transcriptomics, DNA damage response

## Abstract

Space is an extremely hostile environment. Traveling to space has numerous effects on the body of the astronauts at a molecular level. As space agencies prepare for long-term missions, it becomes important to investigate these problems and address them. Thus, in the current study, we assessed the impact of simulated microgravity on the basic pathways especially DNA damage response. We identified a total of 7542 significantly differentially expressed genes. Out of these genes, 4504 were found to be up-regulated while 3038 were down-regulated in the simulated microgravity group in comparison to controls. Pathway enrichment analysis revealed that simulated microgravity has an effect on vital basic biological processes like DNA damage repair, peptide transport, and metabolism. To explore if the same pathways were also altered in humans, we explored the NASA twin study data and found that DDR was also significantly affected in the astronaut but due to ionizing radiation. Upon further investigation, we found that 62 genes belonging to the DDR pathway were mutually differentially expressed in Scott Kelly and the zebrafish embryos. However, there were 29 significantly differentially expressed genes belonging to the DDR pathway in zebrafish embryos that were not found to be differentially expressed in Scott Kelly. Out of these 29 genes, 14 were specific to zebrafish. Upon further investigation, we found that the DDR pathway is affected differently in simulated microgravity as compared to ionizing radiation. These observations provide a preliminary look into the difference in DDR induced by the different factors of spaceflight.

## Introduction

Space travel can have numerous effects on the human body, altering organ and system functions. Some of the major effects have been studied since the Apollo missions, like bone and muscle density loss, nausea, disrupted circadian rhythm, etc^1^. With the advancements in Next-generation sequencing (NGS) technologies, we now have an opportunity to look deeper into the molecular effects of spaceflight^2^. Several studies conducted aboard the International Space Station (ISS), as well as in simulated environments, have shown reversible as well as irreversible effects on various genes and signaling pathways. Functional genomics of astronauts has led to the birth of a new need for personalized medicines for astronauts^3^. Immune dysregulation has been one of the many noticed molecular effects of spaceflights^4^. Several studies have found similarities between the spaceflight-induced immune system and those occurring on Earth^5^.

Spaceflight is a combination of several factors, namely altered gravity, ionizing radiation, the altered concentration of atmospheric gases, and many more^6^. Unfortunately, it is not always possible to conduct a comprehensive study on the effects of all these spaceflight factors on humans as both astronauts and space missions are limited in number. To address these challenges and explore how spaceflight factors affect biology as it happens on earth, several model organisms have been sent to space^7^. Model organisms have their pros and cons when conducting experiments aimed at understanding human biology. Since it is not always feasible to send a biological sample to space, several simulated environments have been designed to replicate spaceflight environments^8–11^. Nevertheless, such experiments do provide insights that can be further validated in human studies.

For example, DNA damage response (DDR) is one of the widely studied pathways in medicine. Damage to the DNA can be caused by several factors including radiation and stress. Exposure of single-stranded DNA or generation of double-stranded DNA breaks can lead to a cascade of signaling relays that is known as DDR^12^. The DDR pathway controls several vital pathways like cell cycle transitions, DNA replication, DNA repair, and apoptosis13. The main regulators of these pathways are phosphoinositide 3-kinase (PI3K)-related protein kinases (PIKKs), including ataxia-telangiectasia mutated (ATM) and ATM and RAD3-related (ATR)^13^. DDR is studied because it is fundamental to the initiation of most tumors^14^. DNA damage if improperly repaired can lead to the activation, deregulation, and overexpression of oncogenes that lead to cancer^15^. DNA damage has also been associated with accelerated aging^10^. Rapid aging is a commonly seen consequence of space travel. Scientists estimate that the heart, blood vessels, and bones deteriorate ten times faster in space than on Earth. In the twin study conducted by NASA, astronaut Scott Kelly’s telomeres were found to be lengthened during flight and shortened post-flight. Ultimately, he ended up having shorter telomeres than what he had before going to the ISS. Changes in the length of the telomere have been associated with aging. While a shorter telomere tends to make a person susceptible to cardiovascular diseases, a longer telomere is often associated with cancer.

In this study, we wanted to investigate the effects of simulated microgravity on vital biological pathways like DDR. To explore ways in which simulated microgravity affects biological function, we selected a study where zebrafish embryos were exposed to simulated microgravity conditions^17^. Although humans are very different from zebrafish, it was found that over 70% of the genes found in humans are also found in zebrafish^18^. Moreover, zebrafishes also have organs like humans, namely two eyes, a mouth, brain, spinal cord, intestine, pancreas, liver, bile ducts, kidney, esophagus, heart, etc. It was found that several genes and important pathways that are required to grow these features are highly conserved between humans and zebrafish^18^. Therefore, it is safe to assume that the patterns of gene expression changes will be similar in humans and zebrafish. Moreover, DDR in zebrafish embryos has been widely studied. Literature review suggests that XRCC5 is an essential gene in the DDR pathway that encodes for Ku80, an essential component of the non-homologous end-joining pathway of DNA repair^19^. Unrepaired DNA damage can lead to unusual apoptosis in the central nervous system during embryo development. In addition to the zebrafish study, we decided to use a dataset generated by the NASA twin study to validate our findings^20^.

## Methods

### Dataset

In this study, we take the data from the public domain available on the data repository of NASA Gene Labs^17^. 12 wild-type zebrafish embryos (Figure-1) of both sexes, and of 3–12 months of age were exposed to simulated microgravity using rotary cell culture system (RCCS), design approved by NASA^21^.

**Figure 1.**
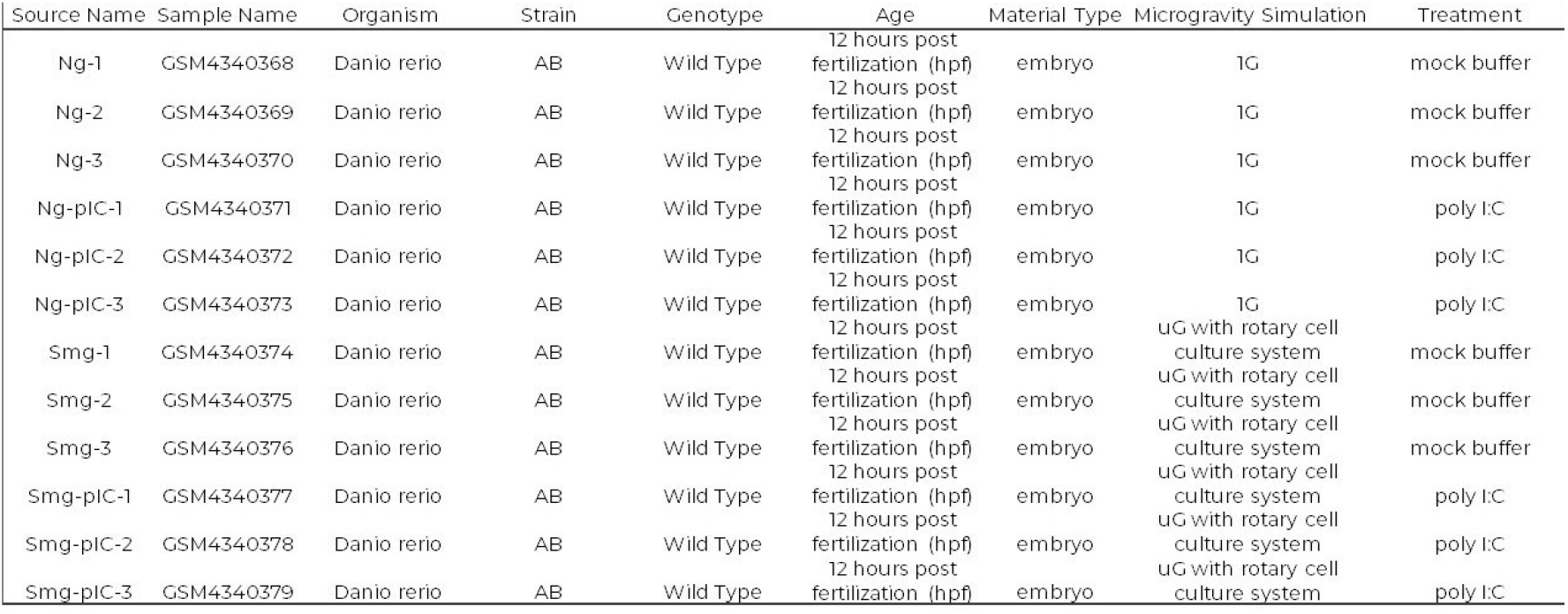
12 zebrafish embryos were classified into four groups, namely Ng – normal gravity without treatment, Ng-pIC – normal gravity with poly-I:C treatment, Smg – simulated microgravity without treatment, and Smg-pIC – simulated microgravity with poly-I:C treatment.

The temperature of 28.5°C was maintained for the duration of the experiments. 12 hours after fertilization, the authors extracted total RNA followed by library construction and RNA sequencing. The reads were amplified in situ. Image analysis and base calling were done using Solexa pipeline v1.8. The quality of sequencing was examined using the standard FastQC software. The adapter reads were trimmed using cutadapt and were aligned to the reference genome of *Danio rerio* (GRCz10) using Hisat2 software. The transcript abundance tables for each sample were estimated with StringTie (v1.2.3), and the Fragments Per Kilobase Million (FPKM) value for gene and transcript level were calculated with R package Ballgown (v2.6.0). The raw gene count generated by the authors was downloaded from NASA Gene Lab^22^ (https://genelab-data.ndc.nasa.gov/genelab/accession/GLDS-373/). The gene count data was log normalized for further analyses. The methods used in the downstream analysis are as follows:

### Exploratory Data Analysis

We performed exploratory data analysis using the PCA (principal component analysis) to understand the patterns in the data. The FPKM gene counts were used for principal component analysis making a scatter plot using R libraries “ggplot” and “ggpubr” on the interactive platform iDEP (v0.94)^23^ (http://bioinformatics.sdstate.edu/idep/).

### Differential gene expression analysis

Differential gene expression analysis was performed using the R package: DESeq2 (https://bioconductor.org/packages/release/bioc/html/DESeq2.html) on the iDEP platform. The two contrasting groups were samples in simulated microgravity (test) and samples in normal gravity (control). The obtained result (a list of differentially expressed genes in samples under microgravity as compared to samples under normal gravity) was filtered by the following parameters: p-value adjusted for False Discovery Rate (FDR) < 0.05 and log2 Fold Change (log2FC) > 2 or <-2 for up and down-regulated genes respectively.

### Pathway Enrichment Analysis

Pathway enrichment analysis was performed using Gene Set Enrichment Analysis (GSEA) and the KEGG pathways database on the iDEP platform. The gene set size used was 15 – 2000 with a pathway significance cut-off (p-value adjusted for FDR) less than 0.05.

## Results

### Exploratory Data Analysis

To understand the overall variation between samples, PCA was performed on log normalized FPKM values. The PCA plot shows a variance of 90% (PC1 + PC2 combined) among samples that are subjected to microgravity as compared to the ones in normal gravity conditions (Figure-2a). It can be seen that the samples that were under normal gravity conditions grouped together. Hence, in downstream analysis, we performed differential gene expression analysis between these two groups.

**Figure 2.**
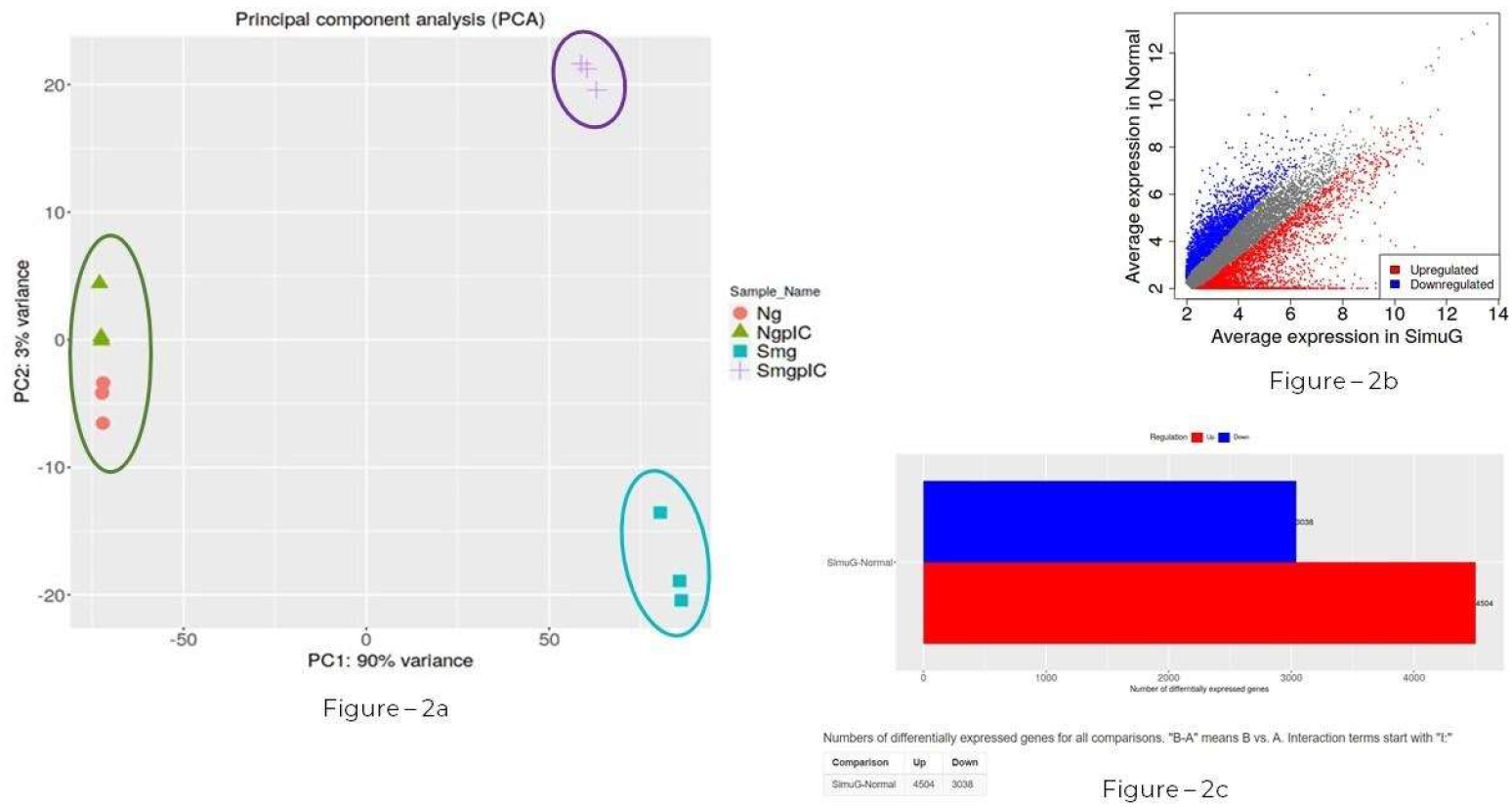
(a) PCA plot showing variance [PC1 (90%) and PC2 (3%)] between zebrafish samples under simulated microgravity and samples under normal gravity (control). Legend: Ng - samples cultured under normal gravity without treatment, NgpIC - samples cultured under normal gravity with treatment, Smg - samples cultured under simulated microgravity without treatment, SmgpIC - samples cultured under simulated microgravity with treatment; (b)– Volcano plot showing the genes that are upregulated and downregulated with average expression in normal plotted in Y-axis and average expression in simulated microgravity plotted in X-axis; (c) 7542 genes were significantly differentially expressed between the two groups. 4504 genes were upregulated and 3038 were downregulated.

### Differential Gene Expression Analysis

Differential gene expression analysis was conducted using DESeq2 between samples under simulated and normal gravity conditions. As a result, we identified 7542 to be significantly differentially expressed (padj. value <0.05, log2 Fold Change ±2) (Figure – 2b). Among them, 4504 were found to be up-regulated while 3038 were down-regulated (Figure – 2c).

To further understand the biological importance of these significantly differentially expressed genes, gene enrichment analysis was performed for all the significantly differentially expressed genes. Gene Set Enrichment Analysis (GSEA) using KEGG pathway gene sets were used to annotate up and down-regulated genes into specific biological pathways (Figure 3).

**Figure 3.**
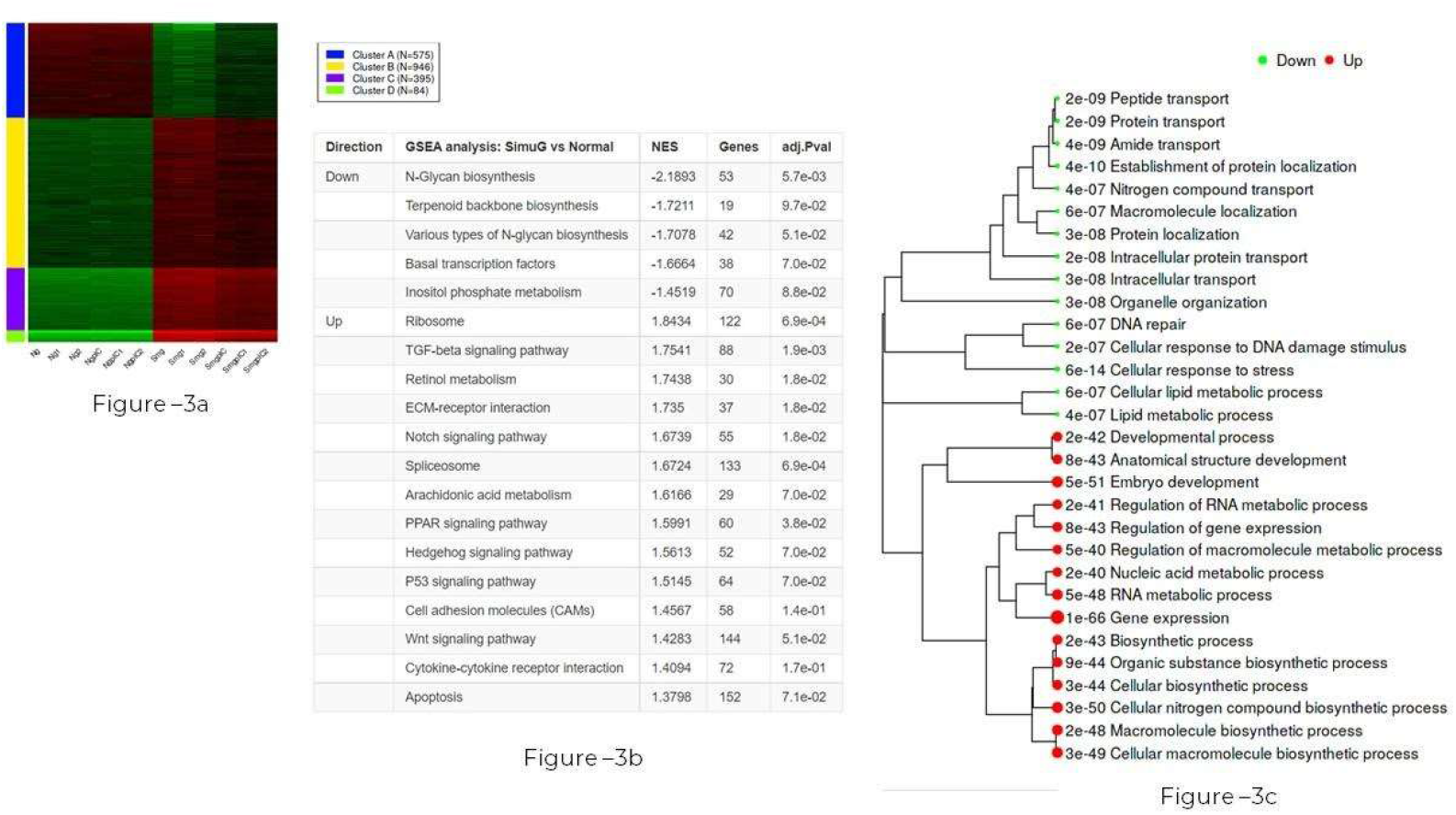
(a) - The heatmap shows the differentially expressed genes divided into four different clusters; (b) – The top pathways that were upregulated and downregulated include several important pathways like Notch, Hedgehog, p53, WNT signaling, and apoptosis; (c)– The dendrogram shows the different pathway clusters and the pathways they include.

The top five most significantly affected pathways were gene expression, embryo development, cellular nitrogen compound biosynthetic process, cellular macromolecule biosynthetic process, and macromolecule biosynthetic process. Cellular response to DNA damage stimulus was also significantly downregulated in microgravity with a padj value of 2.17E-07. The entire list of the pathways can be found in <u>Table 1</u>.

### DNA damage response

DNA damage response works closely along with two other pathways, namely DNA repair and Cellular response to stress. DNA repair pathway consists of 73 genes, Cellular response to DNA damage stimulus consists of 91 genes, and Cellular response to stress consists of 173 genes. Upon comparison, it was seen that 73 genes were shared among all the three pathways whereas 19 genes were common only to Cellular response to DNA damage stimulus and Cellular response to stress pathways (Figure 4). It was interesting to note that 80 genes were unique to the Cellular response to stress pathway. The complete list of genes can be found in <u>Table 2</u>.

**Figure 4.**
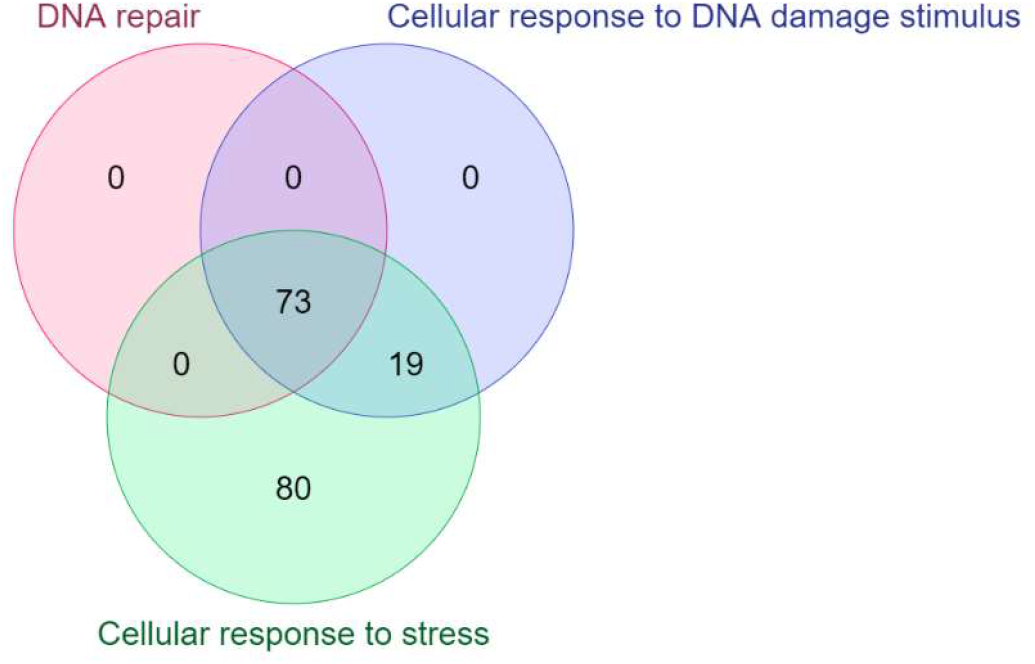
The Venn diagram shows that 73 genes are shared among all three pathways. But only 19 genes are shared between Cellular response to DNA damage stimulus and Cellular response to stress. Interestingly, 80 genes are unique to the Cellular response to stress.

We further went on to compare the pattern of DNA damage response (DDR) pathway in our study with that of the NASA twin study. We found that 62 genes belonging to the DDR pathway were both significantly enriched by differentially expressed genes identified in the twin study^20^ (Scott Kelly) and the zebrafish embryos (<u>Table 3</u>). However, there were 29 significantly differentially expressed genes belonging to the DNA damage pathway in zebrafish embryos that were not found to be differentially expressed in the NASA twin study. Out of these 29 genes, 14 were specific to zebrafish. The remaining 15 genes were identified and listed in <u>Table 3</u>. The authors of the NASA twin study reported that the DDR in the astronaut Scott Kelly was induced by ionizing radiation. In accordance with the genes found in the literature review, we found that the genes: ATM, ATR, and XRCC5, were differentially expressed in the NASA twin study. These are some of the well-known genes to be affected in radiation-induced DDR. However, none of these genes was found to be differentially expressed in the zebrafish embryos. Apparently, there is a difference in the pattern of DDR induced by ionizing radiation and simulated microgravity. Several studies have tried to understand the synergy or interplay of radiation and microgravity in DDR^24^. We propose that simulated microgravity has a different effect on the DDR pathway as compared to radiation. As a result, different gene sets of the DDR are altered in the twin study (induced by ionizing radiation, as claimed by the authors) compared to the zebrafish embryos (induced by simulated microgravity).

Apart from the pathways mentioned above, pathways like Notch signaling, p53 signaling, and apoptosis were also significantly affected in the zebrafish embryos.

## Discussion

Spaceflight, an amalgamation of several unique factors like microgravity, ionizing radiation, and altered atmospheric pressure, has been known to cause many health issues since human beings traveled to space for the first time^1^. While problems like muscle and bone density loss were identified early on, changes at the molecular level can now be studied in great detail thanks to the affordable and convenient NGS technology^25^. Comparative genomics of organisms under spaceflight against ground control have shown several important genes and pathways to be significantly altered4. Before embarking on long-term missions to Mars and elsewhere, researchers should continue to leverage simulated studies to understand the pathways affected by these factors independently and jointly. Such studies could lead to important mediation approaches to preserve astronaut health and recover after space flight.

Traveling to space is a costly affair. Therefore, to study the effects of spaceflight on organisms at the molecular level, scientists often rely on a simulated environment. Simulated microgravity can be established using various techniques like a clinostat, rotating cell culture system, or magnetic levitation^8^. The major question about such experiments here is the accuracy of observations under such simulated conditions. Spaceflight is a combination of several factors^6^. Thus, an isolated factor is not likely to be an accurate representation of real spaceflight conditions.

As we know, it is not always feasible to experiment on humans. Therefore, the majority of space omics research is done on model organisms. But the question that arises is whether the data that we obtain from such experiments are representative of human data? Upon comparing the differentially expressed genes in our study (zebrafish) with the differentially expressed genes of human astronauts generated by the NASA twin study (Cell type: CD4, CD8, and CD19; and coefficient: spaceflight vs ground), we found 26 genes that were mutually differentially expressed in both cases (<u>Table 4</u>). However, it is interesting to note that some of the genes showed dissimilar expression patterns. There could be many explanations for such differences, including the difference in age of the samples studied, the difference in the time duration of the experiments, or the hypothesis that simulated microgravity is not an accurate mimic of real spaceflight. Whatever the reason might be, it should be investigated further to determine how such simulations can be useful to address astronaut health questions.

Omics technologies are fundamental for biomedical research and innovation in human health, food and nutrition, drugs, agriculture, and ecology research^26^. One of the promising areas of omics research is “space omics” which utilizes the understanding of omics to investigate the effects of spaceflight on the genome, transcriptome, metagenome, epigenome, proteome, metabolome, and phenome of the astronauts along with integrative multi-omics approaches. Multi-omics analysis of astronaut omics data and model organisms flown in space provides insight into fundamental biological mechanisms affected by spaceflight^27^. With the establishment of a dedicated space omics data repository and analysis platform: NASA Gene Labs, it has become possible to mine such data. The platform offers easy access and analyses of data generated from real spaceflight missions and simulated experiments^28^. Also, with the formation of an International Consortium for space omics (ISSOP) (https://issop.space/), scientists and students from all over the world can now participate in space omics research projects^29^. With the help of omics, we can better analyze our neighboring planet Mars and look for signs for potential microbial life. Omics will also be important to ensure the safety of the astronauts who will travel to Mars in the near future. Mars is different from Earth in many ways, the most basic one being gravity. This might call for a slightly different approach to omics research on Mars, which would be termed as “Mars Omics”^30^.

## Conclusion

Simulated microgravity has an impact on numerous biological pathways in living organisms. In the study of zebrafish embryos, pathways like DNA damage response and peptide transport were downregulated whereas RNA metabolism was found to be upregulated. Several independent studies have shown that spaceflight affects the DDR pathway in humans as well. However, it has also been noticed that isolated factors like ionizing radiation and microgravity have different effects on DNA damage response^31–34^. It has also been reported that microgravity might play a role in the repair of DNA damaged by radiation34. This project highlights the need to investigate further the impact of microgravity, radiation as well as other factors of space flight on DNA damage response during spaceflights. DNA damage response is an important pathway because improper repair of DNA can lead to several major diseases, including cancer. All of these could be the result of improper DNA repair and DNA damage response. These pathways can have severe consequences on the health of astronauts who spend around 300 days aboard the International Space Station. These problems also need to be addressed before sending humans to Mars or the Moon for a much longer mission.

## Acknowledgment

This research was performed within the framework of the Omics Logic research fellowship program, supported by Pine Biotech. Data processing and visualization pipelines were run using open-source algorithms available on the platform: http://bioinformatics.sdstate.edu/idep/. I would like to thank Mr. Elia Brodsky and Dr. Harpreet Kaur for their expert guidance and mentoring. I would also like to thank Dr. Aleksey Komissarov for his comments and advice.

